# Deep Brain Stimulation for Parkinson’s Disease Induces Spontaneous Cortical Hypersynchrony In Extended Motor and Cognitive Networks

**DOI:** 10.1101/2021.02.23.432542

**Authors:** Maxwell B Wang, Matthew J Boring, Michael J Ward, R Mark Richardson, Avniel Singh Ghuman

## Abstract

The mechanism of action of deep brain stimulation (DBS) for Parkinson’s disease remains unclear. Studies have shown that DBS decreases pathological beta hypersynchrony between the basal ganglia and motor cortex. However, little is known about DBS’s effects on long range corticocortical synchronization. Here, we use machine learning combined with spectral graph theory to compare resting-state cortical connectivity between the off and on-stimulation states and compare these differences to healthy controls. We found that turning DBS on increased high beta and gamma band coherence in a cortical circuit spanning the motor, occipitoparietal, middle temporal, and prefrontal cortices. We found no significant difference between DBS-off and controls in this network with multivariate pattern classification showing that the brain connectivity pattern in control subjects is more like those during DBS-off than DBS-on. These results show that therapeutic DBS increases spontaneous high beta-gamma synchrony in a network that couples motor areas to broader cognitive systems.

## Introduction

Parkinson’s disease is a movement and cognitive disorder characterized by the progressive degeneration of nigrostriatal dopaminergic neurons. While traditionally treated with dopaminergic medications, when pharmaceuticals no longer provide consistent efficacy or lead to severe dyskinesias, high frequency deep brain stimulation (DBS) of the sensorimotor territory of the subthalamic nucleus (STN) or internal globus pallidus (GPi) has been established as the most effective means of managing the symptoms of Parkinson’s disease (Benabid, Chabardes, Mitrofanis, & Pollak, 2009; Deuschl et al., 2006; Limousin et al., 1995; Schuepbach et al., 2013). The therapeutic mechanism of action, however, is still elusive and poorly understood, in part due to the difficulty of conducting neuroimaging studies in the presence of DBS stimulator hardware, due to artifacts and potential safety concerns with fMRI (Alhourani et al., 2015; Boring et al., 2019; Litvak, Florin, Tamás, Groppa, & Muthuraman, 2020). This limited knowledge has become a barrier to improving the efficacy of DBS while minimizing side effects (Alhourani et al., 2015).

Numerous studies have implicated overactive oscillatory synchrony within the basal ganglia, particularly within the beta band (13-30 Hz), as an important pathological feature of untreated Parkinson’s disease (Alhourani et al., 2020; Brown et al., 2001; Hammond, Bergman, & Brown, 2007; Kühn, Kupsch, Schneider, & Brown, 2006). Studies examining interregional interactions using both fMRI and intraoperative recordings have demonstrated abnormal basal ganglia-motor coupling in Parkinson’s disease (Baudrexel et al., 2011; De Hemptinne et al., 2013; Shimamoto et al., 2013). Network analyses have shown that brain networks become less organized and less topologically efficient as Parkinson’s disease progresses (Olde Dubbelink et al., 2013). Beta band hypersynchrony has also been observed in essential tremor, indicating its importance across other movement disorders (Kondylis et al., 2016; Lipski et al., 2017).

Studies comparing neural response when DBS is on to when DBS is off are critical to relate this hypersynchrony to DBS’s downstream neural effects and therapeutic benefits. Effective stimulation has been shown to decrease beta band hypersynchrony in the basal ganglia, particularly within the high beta band region (21-30 Hz) (Bronte-Stewart et al., 2009; Eusebio et al., 2011). De Hemptinne et al. (2015) used electrocorticography recordings in patients with Parkinson’s disease to show that STN DBS reduces beta phase-amplitude coupling in the primary motor cortex, in conjunction with reducing motor symptoms. Oswal et al. (2016) used magnetoencephalography in conjunction with STN recordings 3-6 days after surgery, while DBS leads were still externalized, to demonstrate that acutely after surgery STN DBS modulates connectivity between the basal ganglia and mesial premotor regions in the high beta band range, though the magnitude of this connectivity modulation was not correlated with treatment efficacy.

But how do these results generalize to outside the basal ganglia and motor cortex? Chen et al. (2020) used invasive electrophysiology to show that stimulation of the STN could identify a monosynaptic connection with the prefrontal lobe that was associated with stopping-related activity. A meta-analytic study of fMRI and PET studies in Manes et al. (2014) showed that both the STN and GPi were coactivated with the inferior frontal gyrus.

A critical question for understanding the mechanism of DBS is how does long range cortical to cortical synchronization differ when stimulation is turned on and do these changes normalize prior Parkinson’s-related abnormalities or introduce new transformations? Here we investigated how DBS influences functional connectivity across cortical regions not accessible in DBS surgery, utilizing MEG and a network compression model based upon spectral graph theory. We hypothesized that DBS increases cortical connectivity, similar to dopaminergic replacement therapy (Stoffers, Bosboom, Wolters, Stam, & Berendse, 2008).

To test this hypothesis, we compared resting-state, whole cortex functional connectivity using MEG in the absence of DBS stimulation to recordings obtained during clinically effective high frequency stimulation. We used data-driven analyses, multivariate machine learning methods, and spectral graph theory approaches to assess network level differences between DBS-on and DBS-off across all frequencies and between all pairs of brain regions (e.g. not restricted to somatomotor networks) in an unbiased manner. In addition, we compared these results using the same methods to age matched healthy control subjects to assess whether differences in functional connectivity in the DBS-off condition compared to DBS-on represented a normalization of functional connectivity. These data driven methods have the disadvantage of being relatively less sensitive to small differences between conditions and groups, but have the advantage of casting a wide net to catch large effects in a statistically rigorous and unbiased manner that can seed additional future hypothesis testing. Our results suggest that turning DBS on increases high beta and gamma band synchrony (26 to 50 Hz) across a broad cortical circuit that includes both motor and non-motor systems. Furthermore, functional connectivity patterns in the DBS-off condition is more similar to age matched controls compared to the DBS-on condition, suggesting that rather than normalization, the increased beta and gamma band synchrony is a result of non-normalizing functional connectivity induced by DBS stimulation.

## Results

### Global Cortical Connectivity Difference

Connectedness at a cortical location is defined as the average phase locking between that location and every point on the cortex. We averaged the phase locking at each frequency to find what frequency bands showed a connectedness difference between the DBS-on and DBS-off conditions, as well as between those conditions and controls (Ghuman, van den Honert, Huppert, Wallace, & Martin, 2017; Gotts, Ramot, Jasmin, & Martin, 2019; Gotts et al., 2012). A significant difference between DBS-on and DBS-off was seen in the high beta/gamma band region from 26 to 50 Hz as shown in Figure 1A (DBS-on greater than DBS-off, p<0.05, cluster-level correction for multiple frequency comparisons). In contrast, DBS-off did not show significant global differences compared to age-matched controls, suggesting that the increased synchrony observed in DBS-on did not reflect normalization of abnormal functional connectivity. When STN and GPi stimulation groups were separated, no significant difference in any frequency band was detected; a larger sample may be required to determine whether there are more subtle differences between STN and GPi stimulation than can be detected in the present study.

**Figure 1.**
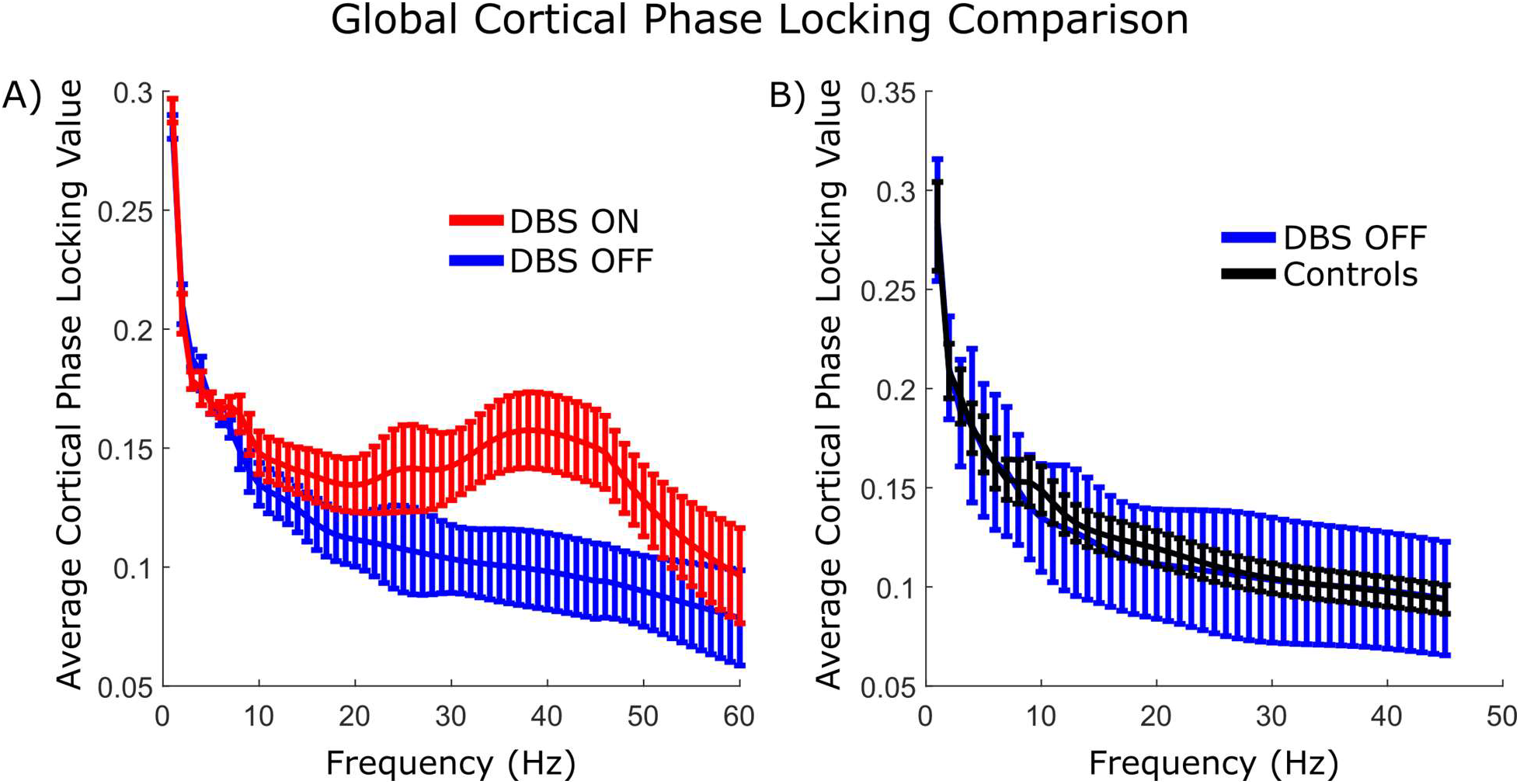
A) Spectral signature of global synchrony when deep brain stimulation is turned on and off. Average phase locking between every pair of cortical points with respect to frequency. Significantly increased beta and gamma band synchrony (26-50Hz) was seen during DBS-on. Error bars indicate paired t-test 95% confidence intervals. B) The spectral signature of healthy controls does not show major deviations compared to the deep brain stimulation off condition. Error bars indicate two sample t-test confidence intervals.

### High Beta Band Networks

All-to-all connectivity networks averaged across the high beta/gamma band (26-50Hz) were computed for each subject for both DBS on and off. To identify a weighted group of connections whose average was consistently changing when DBS was turned on, we utilized spectral graph projections and a support vector machine whose reliability and significance was assessed via crossvalidation. We found that we could identify a pattern of connectivity differences that accurately separated DBS on and off in nine of the eleven subjects (82% leave-one-subject-out cross validated accuracy, p=0.018). Both of the GPi implanted patients were correctly classified, reinforcing that using this relatively broad data-driven analysis, GPi and STN stimulation show similar effects. The largest increases in connectedness occurred in the motor cortex bilaterally, frontal cortex, occipitoparietal lobe, and the right middle temporal gyrus as shown in Figure 2A.

**Figure 2:**
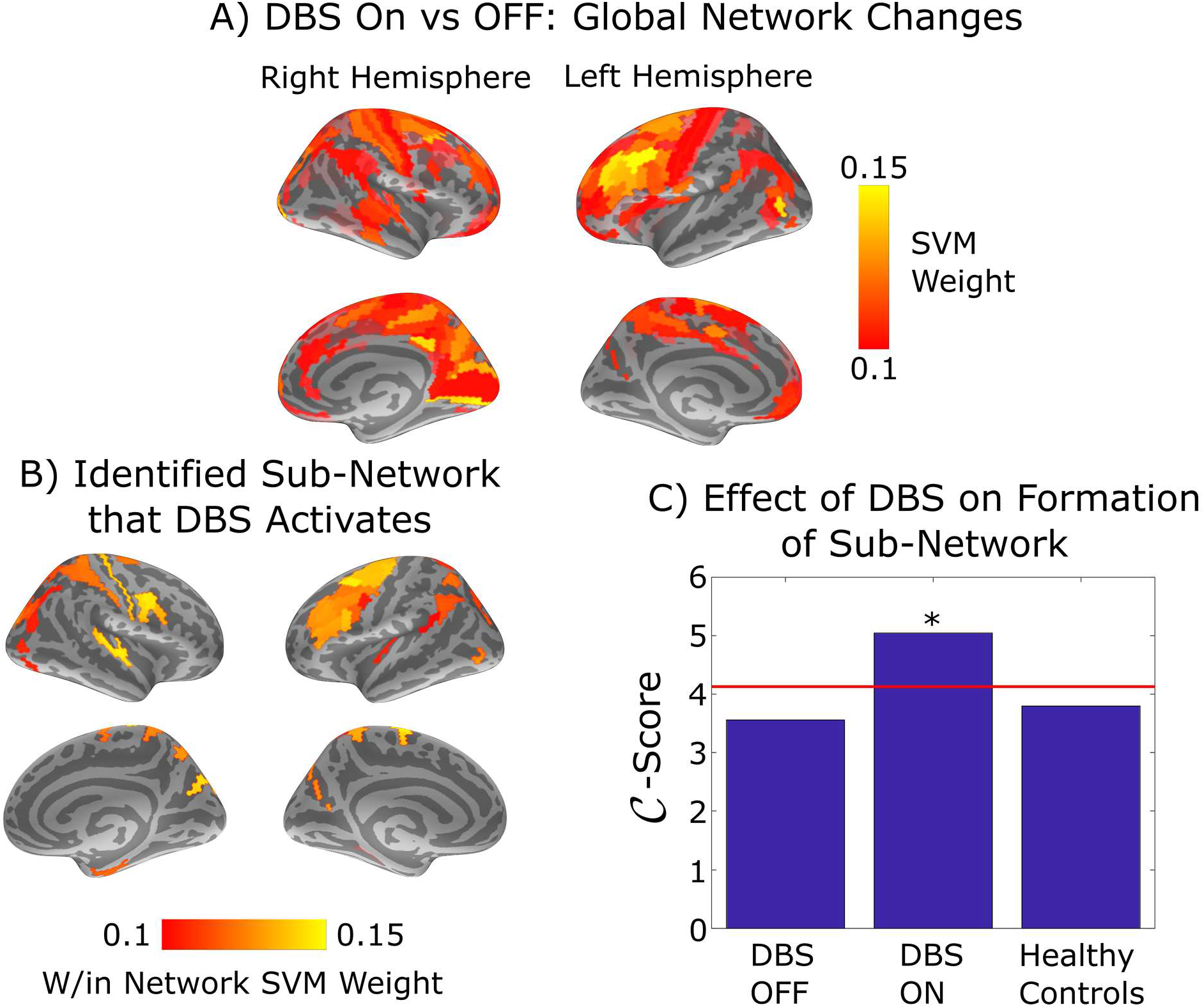
Map of high beta band connectedness. A) Ensemble of connections that were significantly synchronized by deep brain stimulation (DBS) (p=0.018 per cross-validation). Brighter areas indicate larger increases in connectivity with the rest of the cortex when DBS was turned on. B) The connectivity changes from the top figure that forms an inter-connected circuit. A community detection model was used to identify sub-networks whose connectivity within themselves were significantly different across the DBS on and DBS off conditions. Permutation testing revealed one such network, shown here. C) Cluster score of the identified sub-network in the DBS on/off conditions and in healthy controls. The connectivity strength within the subnetwork shown in the bottom-left was compared to strength of equal-sized randomly selected sub-networks to assess whether the identified circuit was significantly activated relative to the rest of the cortex. The red line shows the false detection threshold (α=0.05). The results indicate that discovered circuit’s activation was not significantly distinguishable from the rest of the cortex in healthy controls and when DBS was turned off but was significantly stronger than background when DBS was turned on.

To quantify the relative similarity of DBS-on, DBS-off, and controls, we first used pattern classification to train a model to discriminate the connectivity patterns from the DBS-on and DBS-off conditions and used that model to classify the controls. The resting state connectivity patterns of nearly all controls get classified as being more similar to the DBS-off condition than the DBS-on condition (28/34; p=7.8e-5). Similarly, we trained a model to discriminate the DBS-on connectivity pattern from the control connectivity patterns and used that model to classify the DBS-off data. Classification between controls and DBS-off was not significantly different from chance (48%, p=0.62) and all but one of the DBS-off connectivity patterns were classified as being more similar to controls than the DBS-on condition (10/11; p=0.02). These results show that the connectivity patterns from the DBS-off condition were more like the patterns in controls than in the DBS-on condition.

### Identification of Stimulated Interconnected Circuits

In order to identify interconnected neurological circuits that were being activated by deep brain stimulation, we utilized the Arenas, Fernández, and Gómez (AFG) community detection model. Using permutation testing, the full cortical connectivity changes shown in Figure 2A were clustered into distinct sub-networks (Arenas, Fernandez, & Gomez, 2008). Permutation testing revealed one sub-network that passed statistical significance according to the AFG community detection model, which is illustrated in Figure 2B. This sub-network consisted of four major areas of the cortex: the middle/inferior temporal, occipitoparietal, motor, and the prefrontal cortices.

Figure 2C shows the cluster score for this circuit when DBS is on and off as well as in the healthy controls. Cluster score indicates how well a given sub-network is interconnected within itself relative to rest of the network using a permutation-generated null distribution illustrated by the red line. The circuit illustrated in Figure 2B only emerges as statistically significant when DBS is turned on and is not significant in controls and the DBS off-condition.

### Graph Metrics

Figure 3 illustrates the results of calculating several graph theoretic measures over the found networks and testing to see if the metric changed when DBS was turned on or off using paired t-tests. Global efficiency increased when DBS was turned on, especially within the subnetwork independently identified by the AFG algorithm, indicating that the strongest, most reliable increases in connectivity occurred within that sub-network. The clustering coefficient of the full cortical network was also elevated with DBS, supporting the finding that DBS substantially affects the synchrony of at least one distinct sub-network. There were no significant differences between DBS-off and healthy controls.

**Figure 3:**
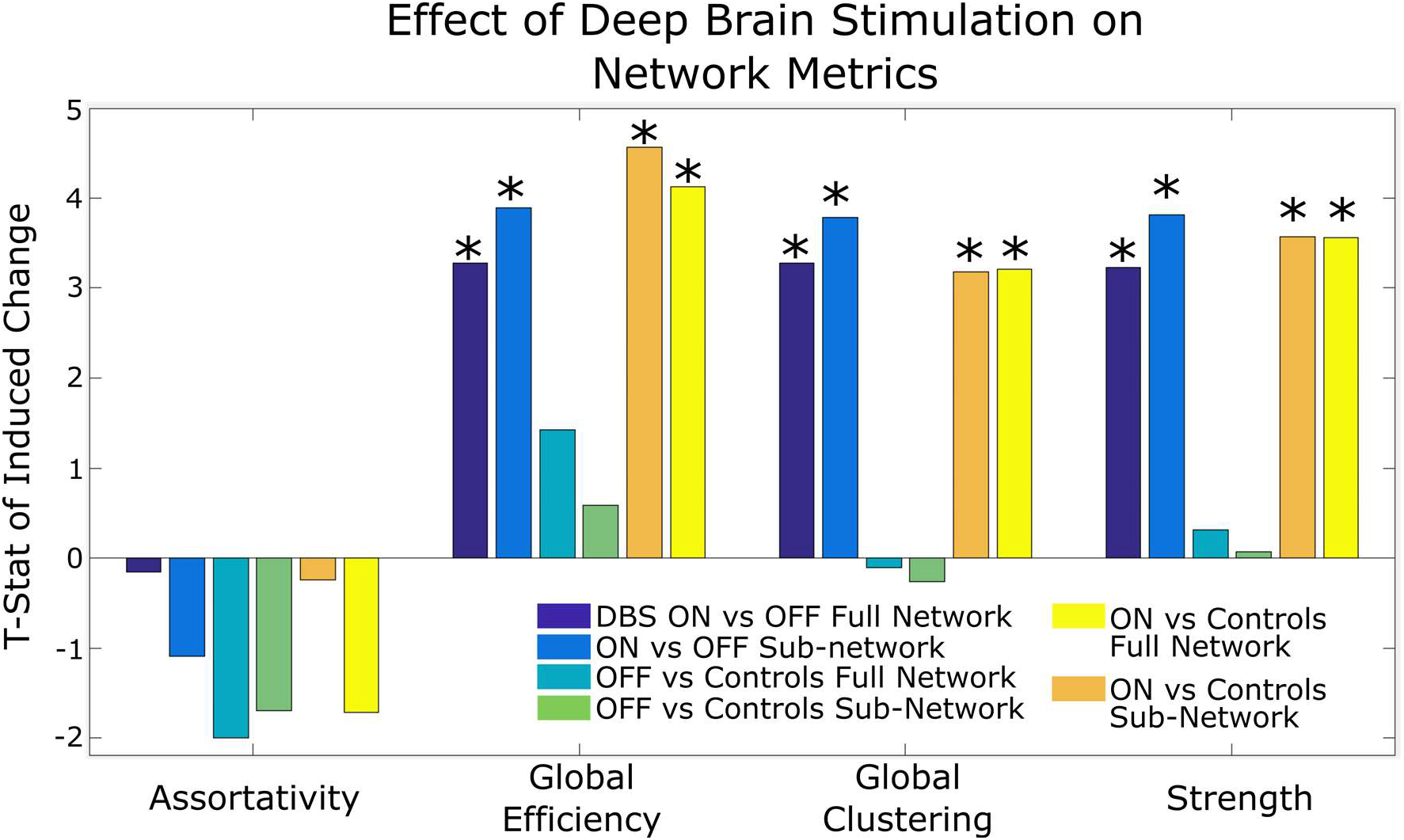
Difference in network metrics between DBS-on and -off using paired t-tests and DBS-on or off vs healthy controls via two-sampled t-tests. These metrics were computed over both the full cortical network (shown in Figure 2A and the identified sub-network (shown in Figure 2B). Positive t-values indicate that the metric increased when DBS was turned on or in DBS-off relative to controls. Assortativity refers to propensity of well-connected brain regions to connect to other similarly well-connected brain regions. Global efficiency is the average inverse shortest path length in the network. Clustering coefficient is the likelihood of regions that are strongly connected to a given region to also be strongly connected to each other. Strength is the average PLV across pairs of dipoles. Asterisks represent statistically significant differences (p<0.05).

## Discussion

We studied the effects of basal ganglia DBS on cortical synchrony in patients with Parkinson’s disease and found that DBS causes increased high beta and gamma band synchrony (26 to 50 Hz). We show that these changes displace cortical networks relative to age-matched controls instead of normalizing them, with these effects being particularly magnified within an interconnected circuit consisting of the motor, occipitoparietal, temporal, and prefrontal cortices. This circuit does not appear to be significantly more activated than the average cortical resting-state synchrony in healthy controls and when DBS is turned off but emerges when DBS is turned on.

### Study Limitations

Several caveats are necessary to consider when interpreting the results of this study. First, we utilized a data-driven approach requiring substantial multiple-comparisons corrections. While this allows us to detect networks that span across non-motor regions that a more targeted approach would not even consider, the tradeoff is that we are only powered to detect very large and straightforward changes. For example, De Hemptinne et al. (2015) found that DBS normalizes coupling locally in the motor cortex between beta phase and broadband amplitude. By focusing on the motor cortex, such a study can pick up interesting changes that our approach is not powered to detect. In general, a lack of detected differences in any category should not be taken as evidence that those differences do not exist.

The second caveat is that the results rely on a sample size of 11 patients and would benefit from validation in a larger cohort, in particular to replicate the non-motor connectivity changes. Third, while DBS was able to effectively control symptoms in the patients utilized in this study, metrics involving relative differences in outcomes were not utilized. Therefore, while the changes in connectivity that we identify can be associated with effective treatment, their association to variability in the degree of individual treatment response would require a more powered study. And lastly, in order to have sufficient power to detect the effects of DBS, we included all subjects with basal ganglia stimulation. When we did separate the GPi and STN stimulation cohorts, neither group was powered sufficiently to detect global cortical connectivity differences. Thus, these results are not meant to represent specific changes resulting from stimulation in either region but rather changes resulting from clinically effective basal ganglia deep brain stimulation.

### DBS modulates long-range cortical connectivity involving the prefrontal cortex, temporal lobe, motor cortex, and occipitoparietal regions

Using our network reduction model, we were able to identify a sub-network of increased cortical connectivity involving the prefrontal cortex, temporal lobe, motor cortex, and the occipitoparietal lobe at the 26-50Hz frequency band. Changes in this frequency band are consistent with the larger literature on Parkinson’s Disease and movement disorders that report aberrant cortical and subcortical oscillations and aberrant basal ganglia-cortical synchrony in this frequency range as being a critical hallmark of these disorders (Brown et al., 2001; Hammond et al., 2007; Kühn et al., 2006).

High beta/gamma band involvement in cortical connectivity differences related to DBS is notable because dopaminergic medication is typically associated with subcortical changes in the low beta band region (12-20Hz) (Hammond et al., 2007; Priori et al., 2004). Furthermore, Bronte-Stewart et al. (2009) also showed that deep brain stimulation of the basal ganglia predominantly attenuates lower beta band power in that region. In contrast, Litvak et al. (2010) demonstrated that increased connectivity between the basal ganglia and premotor areas associated with Parkinson’s occurred mostly in the high beta band. George et al. (2013) also found that dopaminergic medication decreased the number of correlated pairs of scalp EEG pairs mostly at the high beta band (>20Hz). Oswal et al. (2016) showed both properties by demonstrating that DBS decreases basal ganglia power at the low beta band but decreases basal ganglia coherence with the mesial motor cortex in the high beta band. The mechanism of this shift from low beta band synchrony effects subcortically to high beta band synchrony changes in cortical areas may prove an important avenue of future studies, especially in the context of the effects of Parkinson’s and its treatments.

Involvement of the lateral prefrontal cortex, somatosensory, motor/premotor, and occipitoparietal areas are supported by diffusion-tensor-imaging (DTI) and probabilistic tractography findings demonstrating structural connectivity between these regions and the basal ganglia (Lambert et al., 2012; Vanegas-Arroyave et al., 2016). Chen et al. (2020) showed evidence of a monosynaptic STN to prefrontal hyperdirect pathway involved in motor control inhibition, lending further credence to an anatomic basis for this network. The involvement of these regions in Parkinson’s disease and its treatment are also supported by several functional imaging studies (fMRI and PET) (Rowe et al., 2002; Wu et al., 2009). A recent MEG Oswal et al. (2016) study supports the involvement of primary and supplementary motor cortices in the effects of DBS. Connectivity between the temporal lobe and the basal ganglia has been validated by a combination of retrograde transneuronal viral studies and PET studies (Middleton & Strick, 1996; Postuma & Dagher, 2005). Interestingly, Lee, Jang, and Shon (2006) demonstrated that DBS in the basal ganglia was effective in controlling refractory partial epilepsy in patients with temporal lobe epilepsy.

### Effects of DBS displace patients with Parkinson’s relative to healthy controls

In general, we did not find large differences between the DBS off condition and age-matched controls. We do not believe this means they are absent, on the contrary, a large ensemble of literature would indicate the opposite. As mentioned earlier, our sample was most likely not powered enough to detect these differences using a data-driven approach requiring substantial corrections for multiple comparisons. However, the fact that we did see significant differences when DBS was turned on indicates that in contrast to the reported subcortical effects of stimulation, where synchrony is reduced to resemble states observed in subjects without PD, stimulation’s effect cortically appears to be in the opposite direction. A key question for future studies is which of these effects of DBS are associated with therapeutic outcomes, perhaps through compensatory increased synchronization, versus which drive undesired side-effects.

### DBS activated circuit stands out from background synchrony only when DBS is turned on

We found that our DBS-activated circuit’s synchrony was not significantly different from the rest of the cortex in healthy controls and in patients with Parkinson’s when the DBS device was turned off. When the DBS device was turned on, synchrony inside the network increased significantly relative to the rest of the cortex (beyond the overall activation induced by DBS). This increased cortical-cortical high beta synchrony may be a consequence of the release of pathological basal ganglia hyperinhibition seen in Parkinson’s by DBS, leading to the observed network becoming active in DBS-on relative to both DBS-off and controls (Kumar, Cardanobile, Rotter, & Aertsen, 2011; Milosevic et al., 2018). There are two major possibilities for this finding. One is that this cortical network is not typically activated at rest but only during specific tasks, possibly higher-order motor control given the involvement of the premotor cortices. However, when DBS is turned on, this circuit is perturbed as a unit, causing it to also be abnormally activated during resting state. Another is that the magnitude of this circuit’s activation, including at rest, is typically small compared to other networks in the cortex, causing it to disappear into the background of other stronger networks. DBS then causes this circuit to become abnormally active. Further explorations into the state of this circuit under using various stimulation parameters and examining how these effects relate to motor and non-motor behavioral changes with DBS could help mediate between these two hypotheses leading to better understanding of the mechanisms of DBS.

### Conclusions

Studies regarding the effect of DBS in Parkinson’s disease on neural connectivity have largely focused on connectivity within the subcortex and the motor cortex, finding that reduction of overactive oscillatory synchrony, particularly within the beta band, is an important feature of clinically effective high frequency DBS.

We found that DBS introduces new differences in cortical networks of patients with Parkinson’s compared to those from healthy controls in the form of increased connectivity in the high beta and gamma frequency band (26-50 Hz). Most of these changes can be localized to a network that shares several features with that of previously identified cortical motor networks along with the addition of the temporal and occipital regions. Further studies with larger samples are required to link treatment outcomes, and undesirable side effects, to specific aspects of changes in cortical connectivity with DBS shown here. Finding links between particular aspects of neural changes due to DBS and both therapeutic benefit or undesirable side effects could lead to new quantitative paradigms to optimize DBS programming.

## Methods

### Subjects

DBS subjects were eleven patients with bilateral DBS implants for the treatment of Parkinson’s disease, all of whom gave informed consent to participate under STUDY19030378 approved by the University of Pittsburgh Institutional Review Board. Demographic and stimulation information are presented in Table 1. All subjects had implants in either the subthalamic nucleus (STN) or globus pallidus internus (GPi). Stimulation parameters are bilateral unless denoted with left (L) and right (R) designations.

**Table 1:** Patient demographic and stimulation information.

| Age | Gender | Handed-Ness | Location | Stimulation<br>Frequency<br>(Hz) | Voltage (V) | Current (mA) | Pulse<br>Width (μs) | Time<br>between<br>stimulation<br>settings<br>finalized<br>and MEG |
| --- | --- | --- | --- | --- | --- | --- | --- | --- |
| 69 | F | R | STN | 130 | (L) 3.0 (R)<br>11 | 1.70 | 60 | 46 days |
| 71 | M | R | STN | 160 | (L) 3.4 (R)<br>0.0 | 1.70 | 60 | 120 days |
| 72 | F | R | STN | 130 | (L) 2.4 (R)<br>1.8 | (L) 1.47 (R)<br>1.50 | 60 | 91 days |
| 78 | M | R | STN | 160 | (L) 1.0 (R)<br>2.8 | Not recorded | 60 | 71 days |
| 61 | M | R | STN | 180 | (L) 3.3 (R)<br>2.7 | (L) 1.63 (R)<br>1.08 | (L) 60 (R)<br>90 | 388 days |
| 58 | M | L | GPI | 160 | (L) 4.6 (R)<br>4.7 | Not recorded | 60 | 15 days |
| 67 | M | L | STN | 160 | (L) 1.5 (R)<br>1.7 | Not recorded | 60 | 68 days |
| 67 | M | R | STN | 160 | (L) 1.7 (R)<br>1.8 | Not recorded | 60 | 62 days |
| 61 | M | R | STN | 160 | (L) 4.3 (R)<br>1.9 | Not recorded | 60 | 52 days |
| 69 | M | R | GPI | 130 | (L) 2.9 (R)<br>3.0 | Not recorded | 60 | 682 days |
| 58 | M | L | STN | 160 | (L) 4.5 (R)<br>4.7 | Not recorded | 60 | 15 days |

34 healthy controls were selected from a larger population on the basis of age and gender matching. All participants gave informed consent to participate under protocols approved by the University of Pittsburgh Institutional Review Board under STUDY19100015. Healthy controls did not differ in average age (67.8 years with a standard deviation of 5.6 years) compared to the DBS group (66.5±6.3 years, p=0.35). Controls had 21 males, 13 females compared to the 9 males, 2 females in the DBS group (p=0.22).

### Data Collection and preprocessing

Data was collected from 204 gradiometers and 102 magnetometers arranged in orthogonal triplets on an Elekta Neuromag Vectorview MEG system (Elekta Oy, Helsinki, Finland). Data were sampled at 1000 Hz. Electrooculogram and electrocardiogram were concurrently measured to be corrected for during off-line analysis. Head position indicators were used to continuously monitor head position during MEG data acquisition. Signal-space projection (SSP) was performed on MEG data that was subsequently band-pass filtered from 1-70 Hz, notch filtered at 59-61Hz, downsampled to 250 Hz via MNE C scripts, then processed via temporal signal-space separation (tSSS) using a previously validated preprocessing pipeline that cleanses DBS artifacts across DBS-on and DBS-off conditions(Boring et al., 2019). Signal to noise ratio for the inverse calculation was set at nine per Hincapié et al. (2016) demonstrating that higher ratios yielded more accurate detection of changes in connectivity.

Five minutes of resting-state data was collected when the DBS implant was turned on. The implant was then turned off for a half hour, after which another five minutes of resting-state data was collected while the DBS was still off. Resting-state was collected while subjects had their eyes open and fixated on a centrally presented cross. Five minutes of empty room data was also collected. Resting-state data for the controls were collected using an identical protocol.

### Connectivity Analysis

Spontaneous phase locking measures the variability over time of the phase difference between every pairwise cortical location (Lachaux, Rodriguez, Martinerie, & Varela, 1999). We calculated phase-locking values (PLVs) from 1-60Hz and corrected them using empty room noise as described in Ghuman, McDaniel, and Martin (2011). This yielded a 5124 (number of cortical dipoles) x 5124 (number of cortical dipoles) adjacency matrix of pairwise phase locking values between each cortical dipole relative to empty room for each participant at each frequency. To make the data comparable across participants in terms of differential coupling values across frequency bands, we normalized the PLVs with regards to frequency (Schlee, Hartmann, Langguth, & Weisz, 2009). For each participant, we took the distribution of PLVs over all frequencies and calculated their cumulative distribution function and then scaled all phase locking values to this distribution.

### Frequency band selection

To identify a frequency band that displayed significantly different connectivity between deep brain stimulation on and off, we utilized nonparametric cluster level statistics (Maris & Oostenveld, 2007). First, we averaged the PLV across all pairs of dipoles resulting in a 60 (frequency) x 1 vector of the “global connectivity” of a subject’s entire brain network at a given frequency. A paired t-test was calculated at each frequency between DBS on and off and all frequency points with a p-value below 0.05 (not corrected for multiple comparisons as this occurs at the later clustering step) were clustered by frequency adjacency. We then utilized cluster and permutation statistics to find frequency bands that were significantly perturbed by DBS (Maris & Oostenveld, 2007). The connectivity matrices for each subject were then averaged over significant frequency bands to generate a 5124 (cortical dipoles) x 5124 adjacency matrix for each subject. We repeated this protocol except comparing DBS off with health controls. We also repeated this protocol while separating the STN and GPi groups.

### Laplacian Dimensionality Reduction

To identify connections in the cortex that significantly differed between when deep brain stimulation was on and off, we needed to dramatically reduce the dimensionality of the dataset. First we averaged each of the 5124 cortical dipoles across the 360 regions defined in the Human Connectome Project (HCP) atlas (Glasser et al., 2016).

We further reduced dimensionality through an extension of spectral graph theory which states that the critical parts of a network can be understood through a lower dimensional representation utilizing the network Laplacian, a matrix operator intended to reflect the “rate-limiting” steps of a network. This operator has seen increasing usage in the brain connectomics literature as a compact and robust method to analyze both structural and functional brain networks (Abdelnour, Voss, & Raj, 2014; Raj, Kuceyeski, & Weiner, 2012; Wang, Owen, Mukherjee, & Raj, 2017). Here, we studied how the projection of a patient’s connectivity network along these eigenvectors changed when DBS was turned on.

Defining *A*_i,off_ to be the connectivity matrix for *i*-th patient when the DBS electrode is off and *D*_i,off_ to be the diagonal matrix of the degree of each of the 360 regions in the corresponding network. The network Laplacian of the network can then be defined as *L*_i,off_ = *D*_i,off_ – *A*_i,off_. From this we can define an average Laplacian matrix for when the electrode is turned off as 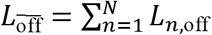. The eigendecomposition of this network would then be 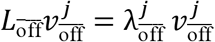 for the *j*-th eigenvector. We can then measure the deviation of a network’s projection along a given eigenvector when DBS is turned on as defined in Equation 1 where 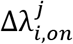 represents the proportional change in the strength of the *i*-th subject’s network when DBS is turned on along the *j*-th eigenvector of the averaged resting-state Laplacian.

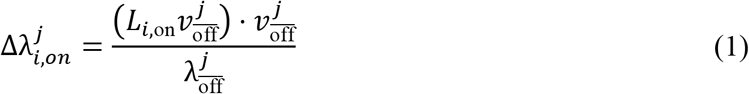

Intuitively, this method of dimensionality reduction can be compared to studying the projection of a dataset along its eigenvector decomposition in principal components analysis. While the eigenvectors in principal components analysis are determined by their ability to capture linear variance in the dataset, the eigenvectors in spectral graph theory are formulated to optimize their ability to preserve several key concepts of a network such as modularity and commute times between nodes (Saerens, Fouss, Yen, & Dupont, 2004). Here, we calculated the projection of each subject’s connectivity matrix along all 360 eigenvectors of the averaged DBS off Laplacian matrix, generating two 360 x 1 feature vectors for each patient: one when DBS was turned on, the other when DBS was turned off.

### DBS On vs Off Classification Algorithm

Our next goal was to identify which particular group of cortical connections were activated by deep brain stimulation. We determined the reliability and significance of our identified ensemble using cross-validation (Browne, 2000).

More specifically, we utilized a support vector machine tested within a leave-one-out crossvalidation architecture. The goal was to present the algorithm with two connectivity profiles, one when the DBS was turned on and the other when it was turned off and have it classify which was which. We accomplished this by formulating two training examples for each subject: one where the 360 x 1 feature vector when DBS was turned on was subtracted from the feature vector when DBS was turned off and the other example the same in reverse. For the algorithm to correctly identify which pair was which, it would have to pick eigenvector components that displayed a large consistent difference between DBS on and off. During cross-validation, both examples associated with the same subject were always placed in the same training fold (e.g. full out-of-sample cross-validation).

The actual algorithm itself consisted of a support vector machine with bootstrapping and random subspace method with parametrization taken from Breiman’s random forest algorithm (Breiman, 2001). To ensure the generalizability of our results, all analyses (frequency band selection, network Laplacian dimensionality reduction, and the classification algorithm training) were performed within a leave-one-out cross-validation training fold.

We repeated this process again for classification between DBS off and healthy controls. For this, we utilized weighted label importance to ensure an even prior for both label classes.

### Identification of Stimulated and Suppressed Communities

We sought to understand whether the changing connections due to DBS self-organized into a specific circuit (sub-network). We utilized the protocol described in (Lancichinetti, Radicchi, & Ramasco, 2010). More specifically, we clustered the change adjacency matrix calculated in Equation 4 according to the Arenas, Fernández and Gómez community detection model (Arenas et al., 2008). The number of clusters was determined according to Newman’s modularity (Newman, 2006). Each cluster was then assigned a 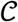-score which detailed how strong the change in connectivity within that sub-network was relative to how strong it would be if the clusters were chosen randomly. The supposition is that a sub-network is considered more significant if connections within it were changing greatly relative to the rest of the network (Lancichinetti et al., 2010).

To generate a null distribution of 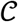-scores, we generated a hundred thousand random undirected, weighted graphs that preserved the edge density distribution of the change adjacency matrix calculated in Equation 4. We repeated the clustering analyses on these random graphs and selected the highest 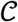-score of the resulting clusters to form our null distribution. For a cluster to be considered statistically significant, its 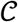-score would have to be within the top five percent of this null distribution.

We also repeated this process on the original resting DBS-off/on and control networks to see whether the identified DBS-activated sub-network was activated significantly prior to DBS and in healthy controls and was simply strengthened by DBS. The permutation process was repeated for each DBS/control group.

We also looked at several traditional graph theoretic metrics within the entire cortical network and the identified significant sub-networks. More specifically, we utilized the metrics outlined in (Rubinov & Sporns, 2010) that were applicable to this situation: assortativity, global efficiency, and global clustering. We calculated these metrics on the original adjacency matrix for each subject both for when DBS was turned on and off utilized paired t-tests to establish statistical significance. We also did two sample t-test comparisons with the healthy controls.

## Acknowledgements

We’d like to thank Ashley Whiteman for her contributions in data collection. This work was supported by the National Institutes of Health (T32GM008208 to M.B.W, T32NS007433-20 to M.J.B., and R01MH107797 to A.G), the National Science Foundation (1734907 to A.G.), and the Brain & Behavior Research Foundation (NARSAD Young Investigator Grant to R.M.R.). M.B.W is also supported by the Hertz Fellowship.

